# Accurate identification of de novo genes in plant genomes using machine learning algorithms

**DOI:** 10.1101/2022.11.01.514720

**Authors:** Claudio Casola, Adekola Owoyemi, Alan E. Pepper, Thomas R. Ioerger

**Author notes:** Author for Correspondence: Claudio Casola, Department of Ecology and Conservation Biology, Texas A&M University, College Station, TX, 77843.

## Abstract

De novo gene birth—the evolution of new protein-coding genes from ancestrally noncoding DNA—is increasingly appreciated as an important source of genetic and phenotypic innovation. However, the frequency and overall biological impact of de novo genes (DNGs) remain controversial. Large-scale surveys of de novo genes are critical to address these issues, but DNG identification represents a persistent challenge due to the lack of standardized protocols and the laborious analyses traditionally used to detect DNGs. Here, we introduced novel approaches to identify de novo genes that rely on Machine Learning Algorithms (MLAs) and are poised to accelerate DNG discovery. We specifically investigated if MLAs developed in one species using known DNGs can accurately predict de novo genes in other genomes. To maximize the applicability of these methods across species, we relied only on DNA and protein sequence features that can be easily obtained from annotation data. Using hundreds of published and newly annotated DNGs from three angiosperms, we trained and tested both Decision Tree (DT) and Neural Network (NN) algorithms. Both MLAs showed high levels of accuracy and recall within-genomes. Although accuracies and recall decreased in cross-species analyses, they remained elevated between evolutionary closely related species. A few training features, including presence of a protein domain and coding probability, held most of the MLAs predictive power. In analyses of all genes from a genome, recall was still elevated. Although false positive rates were relatively high, MLA screenings of whole-genome datasets reduced by up to ten-fold the number of genes to be examined by conventional comparative genomic methods. Thus, a combination of MLAs and traditional strategies can significantly accelerate the accurate discovery of DNG and the annotation in angiosperm genomes.

## Introduction

Novel genes are major drivers of adaptation and evolutionary innovation. A large body of work suggests that new protein-coding genes form at a high rate and represent major contributors to genome evolution and phenotypic variation in plants (1–8). As a result of this rapid evolutionary gene turnover, all species contain hundreds to thousands young, lineage-specific protein-coding sequences that are absent from other taxa (9–12). Both small-scale and whole-genome duplications are responsible for the formation of many novel genes, which tend to share the biological function of their parent genes (1, 3, 4). Occasionally, plants acquired novel genes throughout non-duplicative mechanisms, including horizontal DNA transfer from other species (13, 14) and via the “exaptation” of the coding regions of transposable elements (15–17). Although fundamentally different in nature, all these processes generate new protein-coding sequences from pre-existing genes.

Evolutionary genomic analyses have revealed that new genes can also emerge from ancestrally noncoding DNA sequences wherein an open reading frame (ORF) and new regulatory sequences originate through ‘enabler substitutions’, typically nucleotide substitutions (18–20). Long considered unlikely evolutionary accidents (Jacob 1977), these so-called de novo genes (hereafter, DNGs) encode for evolutionary novel protein sequences that share no homology with genes from other species. Some DNGs have been shown to bear significant phenotypic impacts, from regulating the mating pathway in budding yeast (21, 22), to producing antifreeze proteins in some fish (23, 24) and affecting human health (25–27).

First unequivocally discovered in *Drosophila* (28, 29), de novo genes have been identified across several other groups of animals (23, 30–33) and are well characterized in some model organisms, particularly *Saccharomyces cerevisiae* (21, 34–38). In plants, thousands of potential DNGs have been retrieved through computational surveys in *Arabidopsis thaliana, Brassica rapa*, poplar, rice, sweet orange and *Triticeae* (8, 10–12, 39–47).

Despite the growing number of species and genomes analyzed, the current understanding of de novo gene birth and evolution in plants remain severely limited for several reasons. For instance, the identification of DNGs has traditionally relied on comparative genomic strategies that can be computationally demanding, remain difficult to implement on a large scale, and tend to produce many false positives (48). DNG surveys typically include an initial step wherein all genes (or proteins) in a focal species are queried against genes from other species through sequence homology searches, for example using Blast algorithms (49). The pattern of presence/absence of homologous coding sequences of a given gene along a phylogeny of species allows to estimate its approximate time of origin, a procedure known as ‘phylostratigraphy’ (50). Genes that share no homology outside of a given genome according to the phylostratigraphic method should thus be considered species-specific (taxonomically restricted). However, there are significant caveats associated with phylostratigraphy.

First, homology searches can only generate catalogs of all genes that lack homology, also known as ‘orphan genes’, of which DNGs represent only a subset. Rapidly evolving ancestral genes (51, 52), genes derived from exapted transposable elements (11, 53), horizontally transferred genes (53) and genes with alternative coding frame (11, 53) also contribute to the pool of orphan genes. Although some of these processes are thought to occur at much lower rates than de novo gene birth, the proportion of DNGs among orphan genes might be low (54, 55). Discriminating DNGs from other types of orphan genes requires accurate investigation of synteny conservation across species to identify the enabler substitutions that are uniquely associated with de novo gene birth. Substitutions that enable longer ORFs are especially useful but can be observed only by comparing the DNG coding region with the syntenic genomic regions from several other species (18, 19). While critical to the correct identification of DNGs, the search for enabler substitutions is rarely implemented.

Second, it has been shown that homology searches can underestimate the age of some types of genes, i.e. rapidly evolving genes (56, 57), which can directly affect estimates of DNG rate of formation (48).

Third, homology searches against large datasets require extensive computing resources. Although strategies exist to accelerate this analysis (58), a thorough search of the known sequence space remains challenging and is not reproducible over time, given that sequences databases are expanding exponentially.

Additionally, a wide spectrum of strategies and bioinformatic protocols have been applied to the search of de novo genes in plants and in other organisms. The combination of these issues can produce significant discrepancies in estimates of DNGs even within the same species. For instance, the number of DNGs discovered in *A. thaliana* ranges from 364 (11) to 782 (41). The lack of standardized accurate approaches to assess the number of DNGs represents a major challenge to estimates the rate of de novo gene birth and is diminishing our ability to characterize the biological impact of DNG across plant species.

Machine learning algorithms (MLAs) offer a set of approaches with the potential to mitigate the limitations in DNG detection outlined above. MLAs have proven to be powerful methods for learning models from non-linear datasets in a variety of domains, including many applications in genomics and bioinformatics. In fact, MLAs have been developed to annotate genomic features that include protein-coding genes, RNA genes, enhancers, transcription start sites, splice sites and gene function (59–61). To the best of our knowledge, MLAs applied to the detection of de novo genes have not been developed yet. Interestingly, a few studies have explored the ability of MLAs to identify the broader category of orphan genes in plants (62, 63). A variety of MLAs trained and tested on 1,784 orphan genes from *A. thaliana* showed up to 92% accuracy and 95% sensitivity (62), whereas hybrid deep-learning algorithms applied to 1,544 moso bamboo (*Phyllostachys edulis*) orphan genes reached up to 87% balanced accuracy (63). These results suggest that MLAs can achieve high levels of accuracy for orphan gene prediction. However, as discussed above, DNGs represent only a fraction of orphan genes, and are evolutionarily distinct from other types of genes that lack homology across species. Thus, the ability of MLAs to accurately predict DNGs remains untested.

One of the challenges in applying MLAs to learning classifiers for DNGs is the (typically) small number of positive examples compared to the size of the rest of the genome. While current annotations of plant genomes usually contain tens of thousands of genes, only a few hundred genes can be confidently identified as de novo genes for training. Some MLAs can be sensitive to this asymmetry, outputting models with low information content that appear accurate only because the majority of genes are ancestral genes (hereafter, AGs), while being very inaccurate for DNGs. There are various methods that have been proposed for handling this significant class imbalance (64). We show that sub-sampling of AGs as negative examples can be effective in training accurate models for DNGs. However, although the accuracy on balanced testing sets can be high, even a moderate false positive rate (FPR) can lead to many false positive predictions when the classifier is applied to tens-of-thousands of genes in a whole genome. We show that the FPR can be reduced somewhat by adjusting the selection of examples during training, though at the cost of increasing the false negative rate (FNR). However, false positives can also be removed through traditional comparative genomic analyses that allows to detect signatures of de novo gene birth, i.e. lack of homology in other species and presence of enabler substitutions.

The ability to detect de novo genes through machine learning classifiers depends on the presence of features that show different distributions of values between DNGs and AGs, defined here as genes with no recent de novo origin. Studies across eukaryotes indicate that DNGs and AGs exhibit different distributions in multiple features associated with gene and protein sequences. For instance, DNGs tend to be shorter and with fewer exons than most AGs (8, 11, 35, 36). Proteins encoded by DNGs typically have fewer annotated domains and possess more structurally disordered regions than ancestral proteins in some eukaryotes (18, 20, 51, 65).

Leveraging these observations, we sought to develop and test MLAs aimed at discriminating DNGs from ancestral genes using sequence-derived information. MLA were trained using DNA and protein sequence attributes from DNGs and AGs obtained from three plant species. The de novo gene catalogs consisted of 331 putative DNGs from *Arabidopsis thaliana* (41), 175 and 343 DNGs from *Oryza sativa* (8) and 754 novel DNGs from *Brassica rapa*. These species represent evolutionary lineages with different levels of divergence, as *A. thaliana* and *B. rapa* are relatively closely related species belonging to the Brassicaceae family within the dicotyledon (dicots) clade, whereas rice is a much more evolutionary distant species in the monocotyledon (monocots) clade.

Our investigation had the following goals: (1) Assessing the accuracy and recall of different MLA approaches, including decision trees and neural networks, in discriminating DNGs and AGs; (2) Identifying DNA and protein features with high predictive power for detecting DNGs; (3) Assessing the accuracy and recall of MLAs based exclusively on sequence features compared to those incorporating both sequence features and functional genomic data (gene expression levels, translation level, protein-protein interactions, etc.); (4) Determining the predictive ability of MLAs built on data from one species across other taxa; (5) Determining the predictive ability of MLA approaches in detecting DNGs using whole-genome sequence data that include all genes in a species/accession.

## Results and Discussion

### A set of high confidence *A. thaliana*-specific DNGs

A dataset 782 of putative *A. thaliana*-specific DNGs was recently generated by Li et al. (41). These genes were identified using sequence homology searches on a limited set of databases and without validating the de novo status of each gene through synteny conservation with closely related species. To produce a set of high confidence DNGs from the Li et al. (2016) catalogue, we conducted additional homology searches and retained only genes that passed a series of stringent criteria, including conserved synteny with other Brassicaceae and lack of homology with genomes and transcriptomes deposited on NCBI, as described in **Methods**. This approach follows a robust computational framework developed to identify DNGs (58). A total of 298 putative DNGs were excluded as they shared homology with genes in other species (**S1 Table**), leaving 331 high confidence *A. thaliana*-specific DNGs, similarly to the number of DNGs reported by Donoghue et al. (11); however, we could not directly compare our list of DNGs to those identified in this paper, as de novo gene IDs were not provided by Donoghue et al. (11). The remaining 153 putative *A. thaliana* DNGs shared no conserved synteny with other species and were removed from the catalog as the de novo birth pathway could not be determined.

### Identification of high confidence DNGs in *Brassica*

We first analyzed all available *Brassica* genome assemblies to determine gene annotation completeness. Most assemblies showed a high level of completeness according to BUSCO scores (**S3 Table**). We selected *B. rapa* as the primary focal species for a de novo gene survey due to its agricultural importance and due to the extensive genomic and functional data available for this species (66–71). As the main focal genome to identify de novo genes, we selected the recently improved *B. rapa* v3.0 genome assembly from the Chiifu-401-42 genotype (71), which contained 46,248 protein coding annotated genes and showed a slightly higher annotation completeness than other available assemblies. We performed extensive sequence similarity searches to identify putative *Brassica*-specific genes (see **Methods**) and identified 754 candidate *B. rapa* DNGs.

### Sequence and structural features of de novo genes, ancestral genes and their proteins

On average, de novo gene and protein sequences are known to diverge from ancestral genes in several features. To determine if these qualities can be used to discriminate between DNGs and AGs, we compiled 22 sequence features from the 331 DNGs from *A. thaliana*, the 754 DNGs from *B. rapa*, and the recently published catalogs of 175 and 343 DNGs from rice, *Oryza sativa* (8) (**S1-3 Datasets**). These sequence features are straightforward to calculate for any annotated genome, and do not require any additional data collection (such as gene expression by RNAseq or ribosome profiling). The rice DNGs were obtained integrating sequence homology searches with synteny analyses and were therefore considered well-curated datasets comparable to those of *A. thaliana* and *B. rapa*. To the best of our knowledge, these datasets represent the largest comparative catalog of plant de novo genes to date. The same features were also retrieved by all AGs in these three species (**Table 1**).

**Table 1.** Primary sequence features and mean values of de novo and ancestral genes and proteins.

|  | AT<br>DNGs | AT<br>AGs | BR<br>DNGs | BR<br>AGs | OS<br>DNGs-1 | OS<br>DNGs-2 | OS<br>AGs |
| --- | --- | --- | --- | --- | --- | --- | --- |
| # Genes | **331** | 26,423 | **754** | 34,354 | **175** | **343** | 38,405 |
| Gene length (bp) | **376** | 2,269 | **630** | 1,988 | **4,612** | **3,788** | 3,729 |
| CDS length (bp) | **218** | 1,251 | **350** | 1,169 | **418** | **365** | 1157 |
| CDS #exons | **1.4** | 5.3 | **1.9** | 5.4 | **3.0** | **2.7** | 4.7 |
| GC-content | **43** | 44 | **49** | 47 | **58** | **58** | 58 |
| SSRs in CDS (bp) | **22** | 18 | **16** | 11 | **16** | **10** | 16 |
| %SSRs in CDS | **4.1** | 1.3 | **4.2** | 0.9 | **2.8** | **1.9** | 1.8 |
| Gene overlap | **0.0** | 0.1 | **0.5** | 0.1 | **13.1** | **8.7** | 7.9 |
| CAI | **0.76** | 0.77 | **0.77** | 0.79 | **0.79** | **0.79** | 0.81 |
| %Coding | **31** | 98 | **39** | 92 | **44** | **38** | 84 |
| Coding probability | **0.20** | 0.93 | **0.49** | 0.91 | **0.50** | **0.43** | 0.85 |
| Protein domains | **0.02** | 0.85 | **0.02** | 1.15 | **0** | **0.01** | 0.95 |
| %ISD | **0.28** | 0.31 | **0.33** | 0.30 | **0.46** | **0.44** | 0.34 |
| pI | **8.7** | 7.5 | **8.6** | 7.5 | **9.7** | **9.6** | 8.0 |
| #TM helices | **0.15** | 0.78 | **0.09** | 0.60 | **0.06** | **0.05** | 0.51 |
| %w/TM helices | **13** | 25 | **7** | 15 | **5** | **4** | 14 |
| %w/SP | **15** | 15 | **7** | 11 | **5** | **5** | 12 |
| %w/mTP | **5** | 4 | **3** | 3 | **1** | **1** | 2 |
| %w/cTP | **1** | 6 | **2** | 5 | **2** | **2** | 5 |
| %w/luTP | **0** | 0.5 | **0** | 0 | **0** | **0** | 0 |
| %w/NoTP | **78** | 74 | **88** | 80 | **92** | **92** | 81 |
| Kozak score | **-5.44** | -4.99 | **-5.40** | -5.06 | **-5.15** | **-5.37** | -4.93 |

AT: *A. thaliana*.
BR: *B. rapa*.
OS: *O. sativa*.
CDS: coding sequence.
SSRs: simple sequence repeats (microsatellites).
CAI: codon adaptation index.
Coding probability according to CPC2 (72).
%Coding: proportion of genes predicted by CPC2 to be coding.
%ISD: proportion of protein sequence with intrinsic structural disorder.
pI: Isoelectric point.

Along the lines of previous studies (8, 41), all DNGs components tend to be shorter (except for introns in the *O. sativa* DNG sets), especially at the level of the coding region, and contain fewer exons compared to AGs (**Table 1**; **S2A-C Fig**). The GC-content varied more significantly across species than between DNGs and AGs (**Table 1; S2D Fig**). Interestingly, the GC-content distribution peaks at higher values for AGs in *A. thaliana* and for DNGs in *B. rapa* (**S2D Fig**). In rice, the GC-content distribution is bimodal in AG, as previously described (73), with DNG GC values peaking in between (**S2D Fig**). This pattern in the rice genome is mirrored at the level of the codon adaptation index (**S2G Fig**). Additionally, DNG coding regions contain a higher proportion of DNA derived from microsatellites identified by Tandem Repeat Finder. Interestingly, the microsatellite content is lower in the large rice DNG dataset, which includes older DNGs, suggesting that simple repeat content may decrease with time due to substitutions. As expected, the predicted coding potential of DNGs is much lower than in AGs, with much fewer DNGs being labeled ‘coding’ according to the coding potential calculator 2. In agreement with this, the coding adaptation index of DNGs is on average significantly lower than in AGs (**Table 1; S2E-G Fig**). Furthermore, Kozak scores were lower in DNGs compared to AGs, although their distributions largely overlapped between the two types of genes (**Table 1; S2H Fig**).

Protein features included the presence of conserved domains, predicted proportion of intrinsic structural disorder (ISD) residues, isoelectric point, predicted number of transmembrane helices and subcellular localization peptides (**Table 1; S2I-K Fig**). As expected, given the small size of de novo proteins and their recent origin, very few of them contained conserved domains. Similarly, we found a much lower number of transmembrane motifs (TMs) and genes with TMs in de novo proteins than ancestral proteins. This is in contrast with the observation that *S. cerevisiae* de novo ORFs with adaptive potential are enriched for TM domains (74).

Across all species, DNG proteins showed consistently higher isoelectric point (pI), indicating a higher proportion of basic residues, and very few transmembrane domains compared to AG proteins (**Table 1**). Elevated pIs due to a depletion of acid residues have been also found in mammalian orphan proteins (75) and in S. *cerevisiae*-specific translated ORFs (37), but no explanations for these trends have been put forward. We also found that the distribution of pI values is very different between proteins encoded by DNGs and AGs. While in AG proteins the pI values follow an approximate trimodal distribution, isoelectric points in DNG proteins cluster around two peaks around low (~4) and high (~11) values (**S2I Fig**).

Additionally, we observed higher ISD levels in DNGs than AGs only in *B. rapa* and rice (**Table 1; S2K Fig**). This is in agreement with what observed in *Drosophila melanogaster* DNGs (76) and in orphan genes in *D. melanogaster* (77) and *Leishmania* (78). A similar trend was also reported in rodents young genes (65), although follow up studies suggest that this pattern is likely to be an artifact (51, 79). Some authors have shown that high ISD levels are associated with elevated GC content in orphan genes or young genes (80), possibly a result of some young genes overlapping with ancestral genes (51). It is unclear if the modest difference in GC content between DNGs and AGs in *B. rapa* and rice drives their elevated ISD levels.

Overall, DNG proteins also contained fewer localization peptides compared to AG proteins, with the notable exception in *A. thaliana* of both a higher proportion of mitochondrial transit peptides in DNGs vs. AGs, and a comparable number of DNG and AG proteins with a signal peptide (**Table 1**).

We further investigated possible correlation among features in each species (**Fig 1**). As expected, gene length, CDS length and CDS #exons (the number of coding exons) were positively correlated. Similarly, the coding probability and the presence of protein domains increased with gene length and CDS length and, to a smaller extent, with CDS #exons. Longer coding regions are more likely to be predicted as coding by the coding potential calculator (72). Longer genes are also more likely to encode protein domains. In agreement with previous studies (51, 80), we found that GC-content and ISD were positively correlated. Another expected pattern is the negative correlation between ISD and the presence of transmembrane helices, which cannot form in disordered protein regions. The anticorrelation between GC-content and codon adaption index may be due to the paucity of GC-rich codons among the most-used codons in angiosperms. Similar correlations were observed in *B. rapa* and rice (**S3 Fig**).

**Fig 1.**
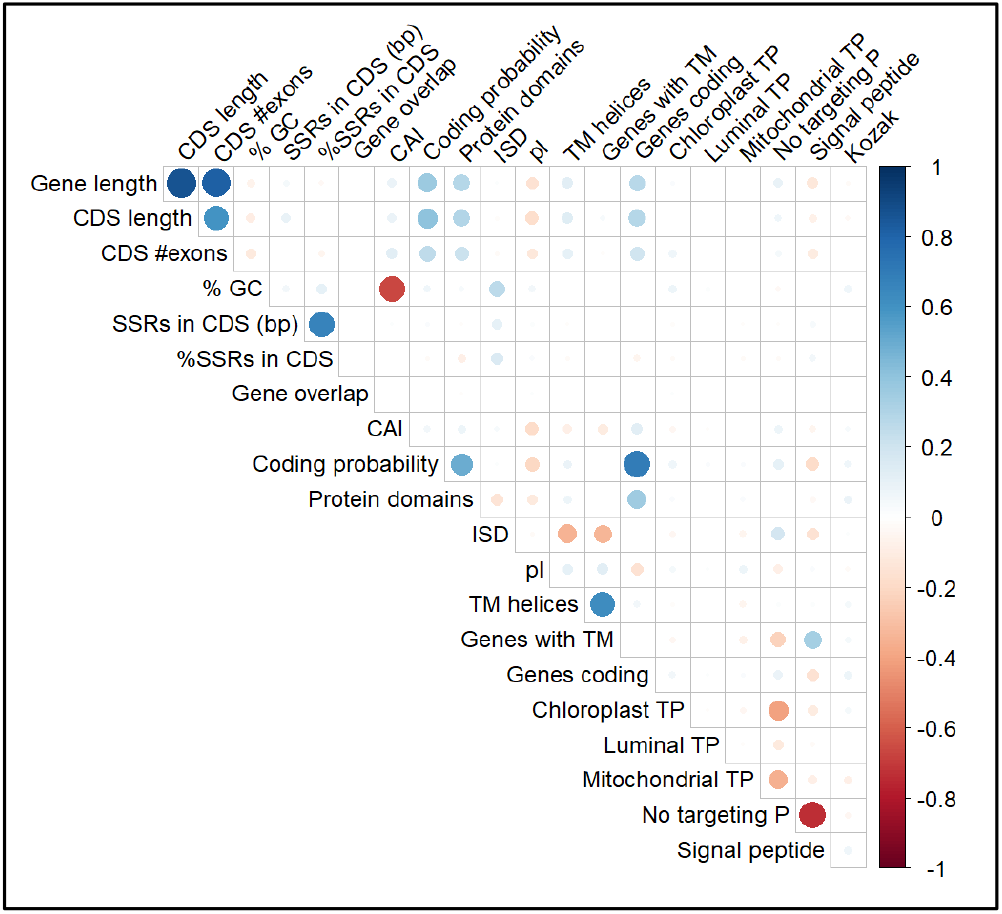
Correlation matrix of *A. thaliana* sequence features used to train ML classifiers. Feature names follow the nomenclature of Table 1.

### Both Decision Tree and Neural Network classifiers detect the majority of de novo genes in test gene sets

As a preliminary attempt to develop a predictive model for DNGs, we trained both a Decision Tree (DT) classifier and a Neural Network (NN) classifier on the *A. thaliana* gene datasets. The set of positive examples was formed by the 331 DNGs we identified in this species. Because using all 26,423 AGs as negative examples led to degenerate tree where every gene was classified as negative due to class imbalance (64), the AGs were sub-sampled (81) by choosing an equal number of 331 genes at random for the negative examples. The NN classifier consisted of a fully-connected network with a hidden layer of 20 units (see **Methods**). We also evaluated the effect of a different number of hidden units (from 10 to 100 in intervals of 10 units) and 2 hidden layers, but these did not significantly improve the accuracy of the NN. When the 662 selected examples were divided randomly into 70% for training and 30% for testing, the DT and NN models were found to have 91-92.0% accuracy in predicting DNGs in the test set. Thus, even though Decision Trees and Neural Networks represent completely different methods for capturing patterns in training data, they are both able to learn how to discriminate DNGs from AGs using sequenced-based features. The confusion matrix for the 100 genes from the 30% test sets shows that the errors are evenly distributed between false positives and false negatives, with recall values above 90% (**Table 2**).

**Table 2.**
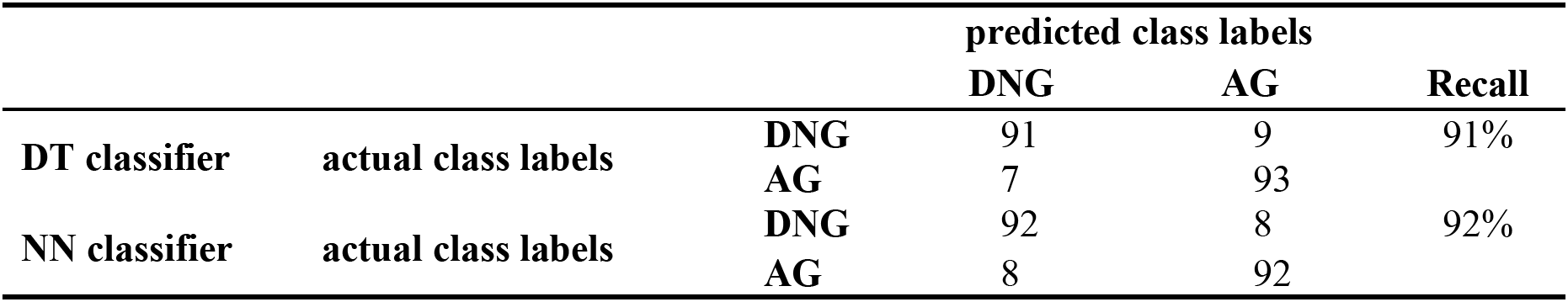
Confusion matrix of DT and NN classifiers, trained on *A. thaliana* genes, applied to an independent balanced test set. Values represent counts of test examples.

|  |  |  | predicted class labels |  |  |
| --- | --- | --- | --- | --- | --- |
|  |  |  | DNG | AG | Recall |
| DT classifier | actual class labels | DNG | 91 | 9 | 91% |
|  |  | AG | 7 | 93 |  |
| NN classifier | actual class labels | DNG | 92 | 8 | 92% |
|  |  | AG | 8 | 92 |  |

In order to determine a more general estimate of the predictive accuracy (since the single tree above is dependent on the specific AGs chosen as negative examples), 10-fold cross-validation was carried out (where a decision tree was generated based on 90% of the data and tested on the remaining 10%, repeated 10 times in a rotated manner), which resulted in a performance estimate of 89.7%, with a 95%-confidence interval (CI95) of 87.3-92.1% (**Table 3**). The same protocol was used to train classifiers on species-specific sequence features retrieved in DNGs and AGs of *B. rapa* and *O. sativa* (using the larger dataset of 343 de novo genes in the latter, see **Table 1**). In all species, NN models performed slightly better than DT models, significantly so in *A. thaliana* and *B. rapa* but not in rice (**Table 3**). DT and NN classifiers showed significantly lower accuracies in *B. rapa* and *O. sativa* compared to *A. thaliana* (**Table 3**; DT *A. thaliana*-*B. rapa P*=0.0038; DT *B. rapa-O. sativa P*=0.0003; NN *A. thaliana-B. rapa P*=0.0005; NN *B. rapa-O. sativa P*=0.0001, unpaired T-test). Overall, these results indicate that MLAs trained on species-specific datasets can successfully retrieve the vast majority of DNGs. Confusion matrices for both classifiers indicate that NN models achieve substantially higher recall than DT models in all species (**S4 Table**). Recall is comparably high (~92%) in NN models of Brassicaceae, but decreased to ~83% in rice, where DT models achieved only ~76% of recall (**S4 Table**).

**Table 3.** Accuracy of DT and NN classifiers in the three angiosperms.

| Species | Decision Tree balanced accuracy<sup>†</sup> (95% C.I.) | Neural Network balanced accuracy<sup>†</sup> (95% C.I.) |
| --- | --- | --- |
| *A. thaliana* | 89.7% (87.3-92.1%) | 93.2% (91.5-94.8)* |
| *B. rapa* | 85.1% (84.3-86.0%) | 88.9% (88.1-89.8)* |
| *O. sativa* | 76.5% (73.1-80.0%) | 80.2% (77.0-83.3) |

<sup>†</sup>Averaged over 10-fold cross-validation
\*NN model vs. DT model within species, *P*-value<0.05, unpaired T-test

The variation in accuracy and recall across species may be due to several factors. A higher quality of gene annotation in *A. thaliana* may explain the increased accuracy of MLAs in this species. The lower accuracy in rice could in part depend on the slightly older age of the larger dataset of 343 DNGs used for this species (8), as DNGs should acquire features that are more of typical genes through time (35). Differences in age of DNGs could also explain the significant overlap in the distribution of continuous features between DNGs and AGs (**S2 Fig**). We also observed that some features show a varying degree of predictive importance between Brassicaceae and rice, which could further contribute to differences in accuracy (see below).

### Training DT classifiers with functional genomic features

We evaluated whether adding functional genomic data could improve the accuracy of the decision tree classifier using datasets available for *A. thaliana*. The classifier was re-trained using 28 additional features which are not systematically available in all genomes, including transcription data (RNAseq), translation levels estimated through Ribosomal profiling (RiboSeq), proximity to transposable elements, selective constraint, and phenotype data for gene knockout mutants (**S5 Table**). The 10-fold cross-validated accuracy of models extended with these functional features was 91.4%, (89.7-93.1%), which is not significantly greater than models without these functional features (*P*>0.05, unpaired T-test). Some of the functional features were occasionally used as decision criteria in lower branches of some of the decision trees; the functional feature with highest importance (0.04) was “AVG RiboP RPKM 25 samples”, which suggests that lack of expression evidence can be an important discriminator for DNGs.

### Feature Importance in Decision Trees

To better assess the contribution of each feature to DT classifiers in the three species, we calculated feature importances, which are based on the relative contribution of each feature to splitting the data in the tree and range between 0 and 1 (see **Methods** and **Fig 2**). Feature importances averaged over 10 runs for each species are shown in **S6 Table**. The DT classifier developed in *A. thaliana* consisted of 70 nodes (including 24 leaves) with a depth of 11. The attribute tested for splitting at the root of the tree, considered the most important based on reduction of Gini impurity, was “Protein domains”. This turns out to be a highly discriminating feature: most AGs (22458/26423=85.0%) have at least one domain (recognized as a known fold family by Pfam based on amino acid sequence), whereas most DNGs do not (only 7/331=2.1% had a recognized domain) (**Fig 2**).

**Fig 2.**
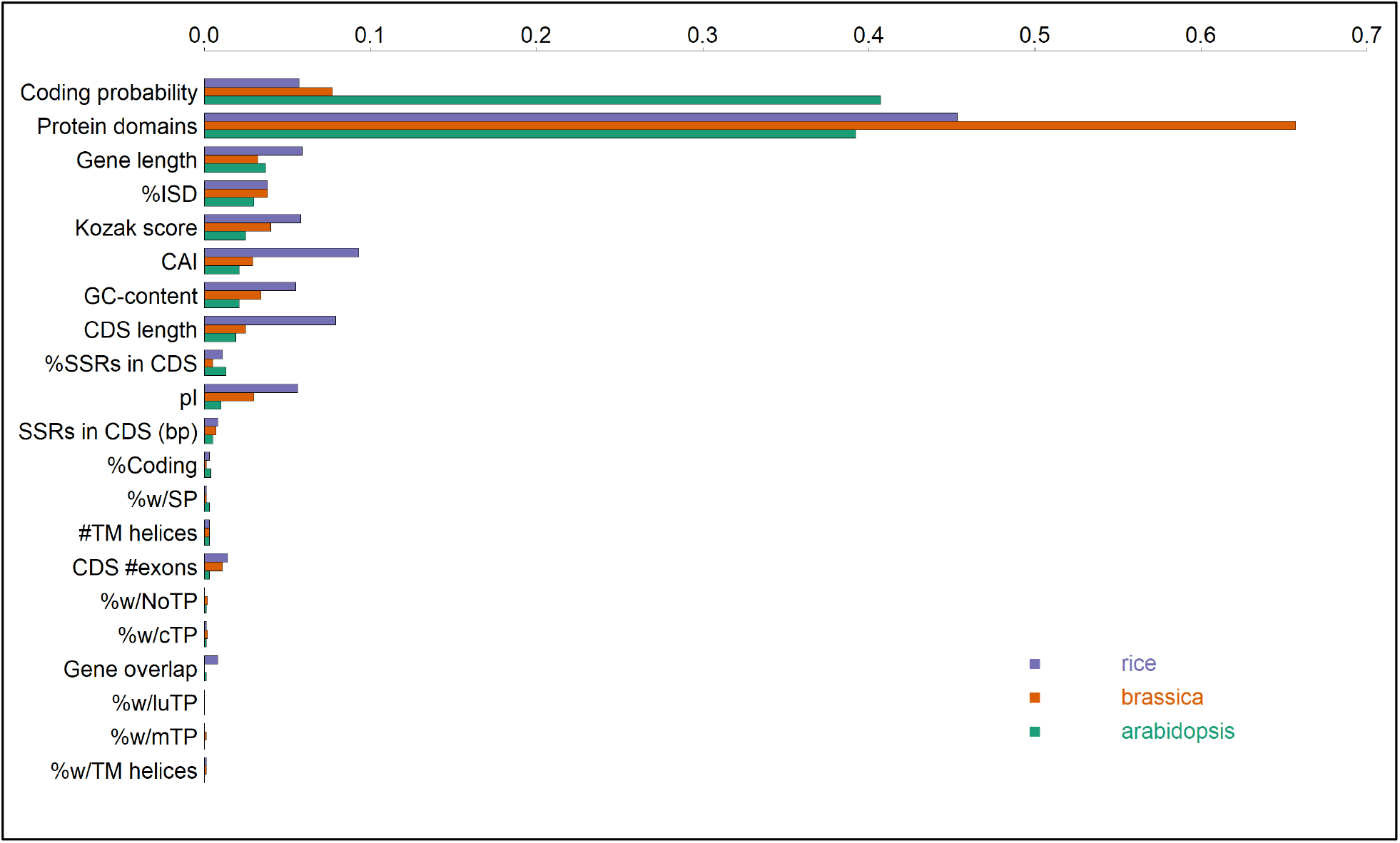
Top ten features in the DT classifier ranked by importance.

Overall, the top ten features by importance are largely the same across all species. “Protein domains” is the most prominent feature across species (**Fig 2**; **S6 Table**). Several other top features that are known to be significantly different between DNGs and AGs, including “Coding probability”, “Gene length”, “CDS length”, “Codon adaptation index (CAI)”, “%GC” and the proportion of “intrinsic structurally disordered (ISD)” regions in proteins, show high importance. Although the presence of a conserved Kozak motif has not been investigated before in DNGs, the “Kozak score” feature showed a relatively high importance, in agreement with other findings suggesting that de novo genes might acquire more ‘gene-like’ regulatory sequences by natural selection after their emergence (31).

Furthermore, some features exhibited substantially higher importance in rice than Brassicaceae. For instance, “Codon adaptation index (CAI)” represents the second most important feature in rice while ranking sixth and eight in *A. thaliana* and *B. rapa*, respectively (**Fig 2; S6 Table**). This is interesting as CAI is only slightly higher in AGs than DNGs in rice (**Table 1; S2G Fig**). “Gene length”, “CDS length”, “Kozak score” are also more prominent features in the monocot.

Interestingly, the top 2 features alone in *A. thaliana* (“Coding probability” and “Protein domains”) can be used to construct decision trees with nearly equivalent accuracy of the ones trained on all features. For example, in *A. thaliana*, the balanced accuracy with such trees (averaged over 10-fold CV) is 91.4% (95%CI: 89.1-93.7%). The decision trees still have multiple nodes in them, typically around 30; they just include splits on multiple threshold values (i.e. sub-ranges) of “Coding probability”. The average feature importances are 0.407 for “Coding probability” and 0.392 for “Protein domains”. In fact, the neural network performs even better, probably due to the reduction in parameters (weights in the network) with just two inputs: 94.0% (95%CI: 92.2-95.7%). Further analyses will be necessary to determine if this applies to other plant genomes.

### Accuracy is not limited by small training set size

The size of the training dataset can affect accuracy in MLA predictions. We tested if this is the case for the DT classifiers using randomly selected subsets of *A. thaliana* de novo genes and ancestral genes equal to 10-100% of the original training set of 464 genes. This was repeated 30 times for each set size to obtain a DT classifier learning curve. The testing dataset for each analysis was carried out on a group of DNGs that did not overlap with the training set. Additionally, during training and testing, the number of negative examples (AGs) was always balanced with an equal number of positive examples. We found that the accuracy of the DT remains elevated (>85%) even for training with one-tenth of the full training-set size of 464 genes and is nearly equal to the highest accuracy with only 30% of the full training size, corresponding to 138 genes (**Fig 3**). The observed trend also suggests that the accuracy would not be significantly improved by using a larger training set with more known DNGs.

**Fig 3.**
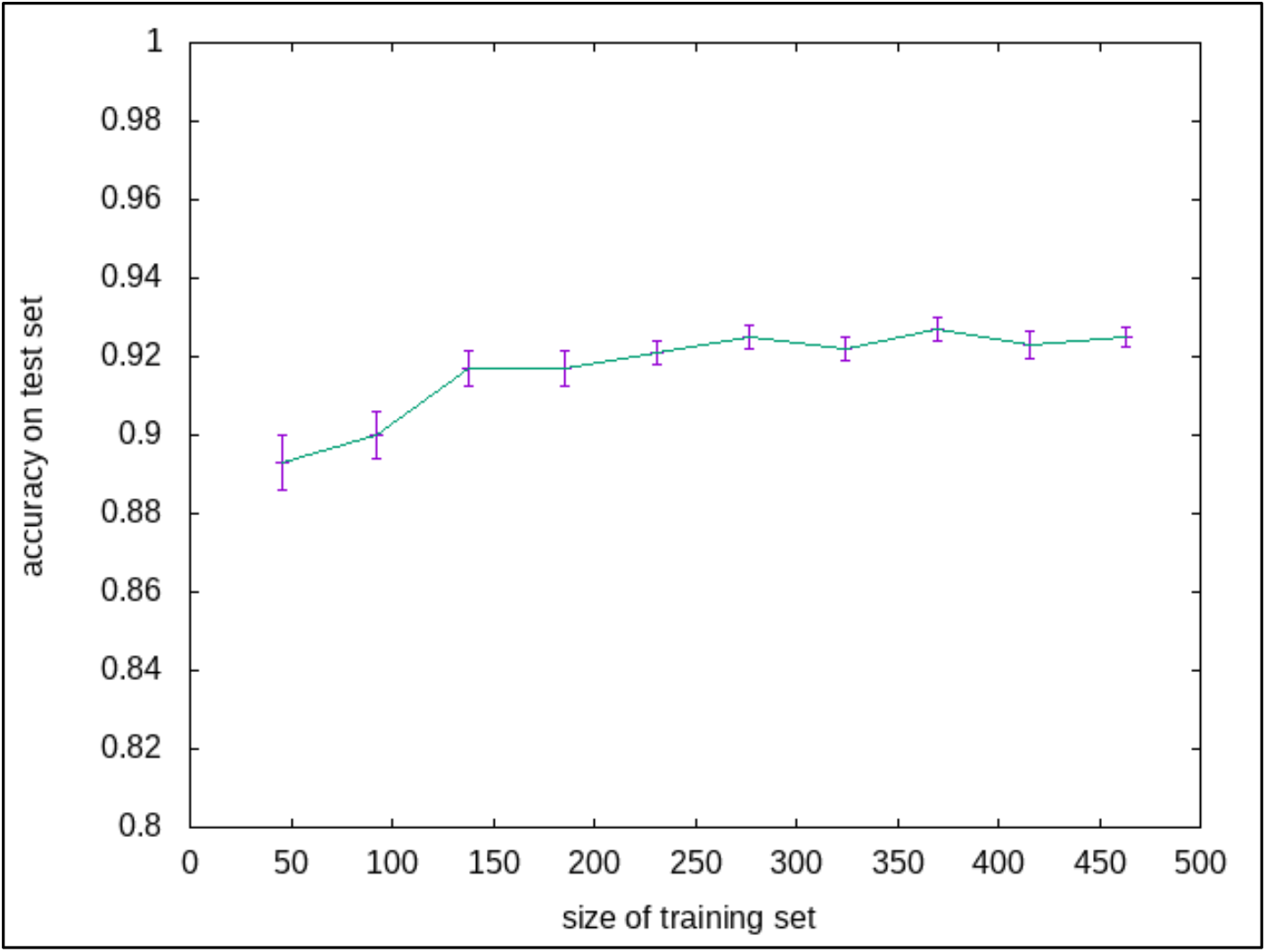
Learning curve of the *A. thaliana* DT classifier on 10-100% of the original training dataset. Standard errors are shown.

### Cross-species models for de novo gene prediction are nearly as accurate as species-specific models

An important question is whether the patterns extracted by the MLAs for discriminating DNGs are species-specific, or whether the MLAs are capturing general properties of DNGs that extend across multiple plant species. To assess this, we trained a DT and NN classifiers using data on genes from *A. thaliana*, and then applied these models to the *B. rapa* and *O. sativa* datasets. In 10-fold cross-validation analyses, the *A. thaliana* DT model achieved 83.1% and 67.2% accuracy in *B. rapa* and rice, respectively (**Table 4**). The *A. thaliana* NN model reached slightly higher accuracy in both species (**Table 4**). Overall, species-specific models (**Table 3**) achieved significantly higher accuracy than cross-species models (*P*-value<0.05, unpaired T-test), except for the *A. thaliana* DT model applied to *B. rapa* datasets (*P*-value=0.0855, unpaired T-test). Taken together, these results indicate that models trained on the *A. thaliana* genome, which has been more carefully and thoroughly annotated, can be applied with nearly equal accuracy on *B. rapa*, and does not require an MLA to be re-trained on each new gene set. Conversely, *A. thaliana* classifiers achieved substantially lower accuracies in *O. sativa* compared to the rice-specific models. It appears that the features associated with DNGs in *A. thaliana*, such as coding probability, lacking recognizable protein domains, and having lower Kozak score, generalize across species, and are associated with DNGs in other plant genomes.

**Table 4.** Comparison of performance of DT and NN models using a common model trained on *A. thaliana* versus species-specific models.

| Species | Decision Tree<br>balanced accuracy<sup>†</sup><br>(95% C.I.) | Neural Network<br>balanced accuracy<sup>†</sup><br>(95% C.I.) |
| --- | --- | --- |
| *B. rapa* | 83.1% (81.2-85.0) | 85.1% (83.5-86.6) |
| *O. sativa* | 67.2% (63.9-70.5) | 69.4% (66.5-72.2) |

<sup>†</sup>Averaged over 10-fold cross-validation

The higher predictive ability of *A. thaliana* MLA classifiers in *B. rapa* compared to rice suggests that cross-species DNG identification with MLAs tend to be more accurate in closely related genomes. Features that differ significantly between Brassicaceae and rice genes, including gene length, %GC and simple repeat content in the coding region, may drive the lower sensitivity of the *A. thaliana* DT classifier in rice. A broader taxonomic sampling at varying phylogenetic distances from *A. thaliana* will be required to test this hypothesis more thoroughly. We noticed that the recall decreased in cross-species prediction, with a limited difference in *B. rapa* (from ~84-92% to ~82-84%) and a significant drop in in rice from ~76-83% to ~55-59%) (**S7 Table**). This indicates that cross-species MLA predictions of DNGs may achieve acceptable levels of sensitivity within taxonomic families but fail to detect a substantial fraction of DNGs in more distantly related species. Thus, broad MLA-based de novo gene surveys in plants may require the training of classifiers using DNGs detected with comparative genomic approaches in at least one species per family.

### Whole-genome predictions of DNGs using DT and NN classifiers

We next assessed the accuracy of species-specific DT and NN classifiers to predict DNGs in whole-genome gene sets. We calculated the balanced accuracy, which is better suited to assess the performance of classifiers when classes are imbalanced. The overall balanced accuracy ranged from 92.3% in *A. thaliana* to 76.9% in *O. sativa*, with slightly higher accuracy in DT vs. NN models (**Table 5**). A high recall of ~94-99% was found across species and classifiers, although DT models also achieved lower FPRs compared to NN models (**S8 Table**). Given the much higher number of AGs than DNGs, the total number of false positives reached ~2,055 in *A. thaliana* and a maximum of ~8,948 genes in *O. sativa* (**S8 Table**). Given the high FPRs, MLAs alone may achieve the level of accuracy required to entirely replace traditional comparative genomic analyses in DNG surveys; however, the application of MLAs as a first step would decrease by up to 10-fold the number of genes that need to be investigated with homology searches and other time-consuming approaches in order to remove false positives.

**Table 5.** Comparison of performance of DT and NN species-specific models applied to whole genomes.

| species | Decision Tree genome-wide accuracy<sup>†</sup> (95% C.I.) | Neural Network genome-wide accuracy<sup>†</sup> (95% C.I.) |
| --- | --- | --- |
| *A. thaliana* | 92.3% (91.9-92.7) | 91.1% (90.4-91.9) |
| *B. rapa* | 87.4% (87.1-87.7) | 86.0% (85.6-86.4) |
| *O. sativa* | 79.3% (78.7-79.9) | 76.9% (74.9-78.8) |

<sup>†</sup>Averaged over 10-fold cross-validation

We further examined if the performance of models trained on *A. thaliana* data but applied to the two other species (in this analysis, there are no confidence intervals because a single input model trained on one species was tested for accuracy on the whole genome of another). The *A. thaliana* classifiers, particularly NN models, showed relatively high <u>balanced</u> accuracy in *B. rapa* and in rice (**Table 6**). However, recall values dropped significantly in both species, from ~97-98% to ~82-84% in *B. rapa* and from ~94-98% to ~55-59% in rice (**S9 Table**). Interestingly, the *A. thaliana* NN model resulted in lower FPRs than the within-species models while the opposite was true for the *A. thaliana* DT model (**S9 Table**).

**Table 6.** Comparison of performance of DT and NN models using a common model trained on *A. thaliana* versus species-specific models applied to whole genomes.

| species | Decision Tree<br>genome-wide accuracy | Neural Network<br>genome-wide accuracy |
| --- | --- | --- |
| *B. rapa* | 84.9% | 86.2% |
| *O. sativa* | 74.6% | 81.6% |

### The DT classifier specificity can be adjusted by increasing the proportion of negative examples during training

Given the high false positive rates in classifiers, we investigated if including negative examples during training could increase the model specificity using *A. thaliana* datasets and the DT model. We scaled up the number of negative examples up to 10 times the original set of 232 AGs while maintaining a balanced test set with equal numbers of DNGs and AGs. Each iteration at different training sizes was repeated 30 times. We found that for increasing numbers of negative examples, the balanced accuracy steadily decreases from ~91% to ~85%, primarily due to the increasing false negative rate from <10% up to 25% (**Fig 4**). Concomitantly, the false positive rate decreased from ~8% to <4% (**Fig 4**). As the main goal of the MLA approach is to detect DNGs, the loss of sensitivity associated with the higher number of negative examples might be not worthwhile. However, we noticed that this tradeoff between false positive and false negative rates might be acceptable for limited increases in the size of negative examples.

**Fig 4.**
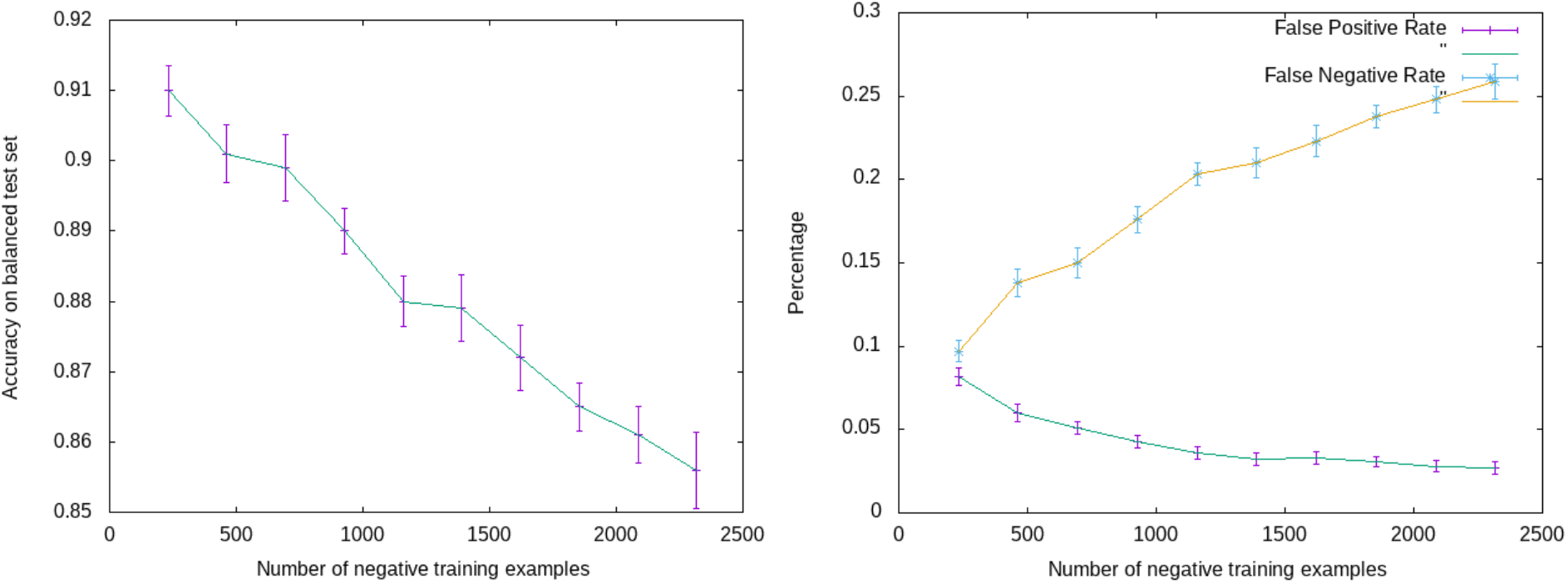
Accuracy (left) and false positive and false negative rates (right) for increasing number of negative examples used in the training of the DT classifier in *A. thaliana*. Standard errors are shown.

## Conclusions

In this study, we have developed and assessed the first machine learning framework to identify de novo genes. In order to make these approaches readily applicable in species with limited functional genomic data, we have specifically selected basic sequence features that can be obtained from DNA and protein sequences available in annotated genomes. Using DNG datasets from three plant species, including an updated gene set from *A. thaliana* and the first group of de novo genes in *B. rapa*, we have found that both decision tree (DT) and neural network (NN) classifiers achieve high levels of accuracy and recall in predicting DNGs. Using DT algorithms applied to sub-sampled sets of DNGs and AGs, we identified a few features with significant predictive power for DNGs. This is in line with performance ability of MLAs to discover orphan genes in *A. thaliana* based on six DNA sequence features (62). Importantly, orphan genes are not equivalent to de novo genes, as the former appear to be mostly constituted by rapidly evolving genes (54). Training MLA models with additional features derived from functional genomic data (transcription and translation data) and information from phenotypic assays that are not readily available in most sequenced genomes does not lead to a substantial increase in accuracy or reduction in the number false positives.

A major advantage offered by MLAs is the significant decrease in computational time compared to traditional genomic approaches to find DNGs—essentially, a time contraction from weeks or months to minutes. For MLAs to be successfully applied in DNG surveys across hundreds to thousands of species, it is critical to train models using datasets of known DNGs from a few key species, and for these models to obtain high accuracy and recall in other species. We conducted initial tests to explore this possibility using DT and NN models trained on *A. thaliana* to predict DNGs in *B. rapa* and rice. We found that these cross-species predictions achieved comparable accuracy of species-specific models in *B. rapa*, and somewhat lower accuracy in rice. This suggests that MLAs trained in one species can likely be used to infer DNGs in closely related species, as in the case of the two Brassicaceae, *A. thaliana* and *B. rapa*. Given that many angiosperm families now contain sequenced genomes from multiple species, and considering both the rapid increased of the number and quality of new genome assemblies, de novo gene discovery based on MLAs could likely be applied to a large number of flowering plant taxa. Future work on more taxa should help better determine how cross-species MLAs accuracy decreases when the evolutionary distance between taxa increases, as this study indicates.

Genome-wide analyses showed that species-specific models predicted well above 90% of known DNGs, although the much higher number of ancestral genes lead to several thousand false positive cases. In cross-species genome-wide analyses, *A. thaliana* models identified 82-84% of true DNGs in *B. rapa*, but achieved less than 60% recall in rice. Notably, FPRs were substantially lower in cross-species NN models compared to species-specific models. The combination of high recall, at least in some cross-species tests, and high FPRs suggests that a three-step pathway can be employed to accelerate DNG discovery in angiosperms. First, NN models are developed in one or two species with the best gene annotation quality in each angiosperm family. Second, these NN models are applied to an array of target species in the same family. Third, the candidate DNGs predicted from each target species, comprising only a few thousand genes, are analyzed post-hoc with traditional comparative genomic approaches to remove false positives.

As this represents the first systematic study to assess machine learning approaches in de novo gene discovery, we expect that further developments of in this area could significantly increase accuracy and recall while reducing the false positive rate in DNG detection. Along these lines, alternative machine learning approaches, including deep neural networks, and methods to address the class imbalance between DNGs and AGs different from sub-sampling, such as synthetic minority over-sampling algorithms, or SMOTE (62, 82), warrant future investigations.

## Methods

### Validation of *Arabidopsis thaliana* de novo genes

The TAIRv10 (TAIR10) DNA and protein sequences of *A. thaliana* were obtained from the folder “TAIR10 blastsets” in the TAIR repository (83). Data from the files “TAIR10_pep_20101214”, “TAIR10_cds_20101214” and “TAIR10_exon_20101028” were used in sequence similarity searches and sequence feature analyses. *A. thaliana* de novo genes were retrieved from a set of 782 putative DNGs recently described by Li et al (2016) using sequence homology searches. These genes were screened to identify high-confidence DNGs supported by further comparative genomic data, particularly synteny information. Specifically, we performed Blast v2.11.0 (49) searches of the corresponding protein sequences against several NCBI databases. We used Blastp to search the “nr” and “tsa_nr” databases, and tBlastn to search the “nr/nt”, “refseq_rna”, “est” and “TSA” databases with the following modified parameters: num_descriptions 100 -num_alignments 100 -max_hsps 5 -evalue 0.001 -seg yes. The seg filter was turned on in order to remove spurious hits due to nonhomologous stretches of similar amino acids. These searches were carried out against increasingly broader taxonomic units that include *Arabidopsis* species excluding *A. thaliana*, Brassicaceae (excluding *Arabidopsis*), rosidae (excluding Brassicaceae), angiosperms (excluding rosidae), green plants (excluding angiosperms). We also searched for homologous sequences of the 782 DNGs in fungi, bacteria and archaea to identify and remove possible horizontal transfer cases (**S1 Table**). Subject sequences were screened using unix scripts to remove truncated proteins. We excluded from the catalog of DNGs all cases with any homology with sequences from a non-focal species.

To determine synteny conservation of DNG coding regions we searched 45 Brassicaceae genome assemblies obtained from Phytozome (https://phytozome-next.jgi.doe.gov) using the translated DNA sequence of each exon of the 782 putative DNGs, which allowed us to detect conserved coding regions for genes with one or multiple exons, in tBlastx run with the following modified parameters: num_descriptions 100 -num_alignments 100 -max_hsps 5 -evalue 0.001 -seg yes. A list of the 45 species investigated in available in **S2 Table**.

First, we used these alignments to identify Brassicaceae genomic regions that were syntenic with DNGs and with the potential to encode proteins. Although these regions did not include annotated genes, they maintained long coding regions and were thus considered *bona fide* homologs to DNGs. We based this selection on two criteria. We selected for hits with a conserved methionine within the first five amino acids of the *A. thaliana* DNG protein, in order to account for alternative first codon positions (84). Additionally, we included only hits wherein the total alignment length from the first codon to the first stop codon equal to at least 75% of the query protein. This threshold was selected to include hits that are likely to encode a protein, given the lack of disabling mutations along most the of the coding, while allowing for slightly shorter loci, as stop codons may also vary slightly between orthologs. Using this strategy, we identified 44 DNGs with putative homologous genes in non-*A. thaliana* Brassicaceae, which were thus discarded.

Second, we used the same alignment data for the remaining DNGs to identify those that maintain synteny conservation *in noncoding regions* with other Brassicaceae. To this end, we applied a minimum threshold of 30% coverage between *A. thaliana* DNGs and Brassicaceae genomes as corresponding to conserved synteny. This length threshold is lower than those used in previous DNG analyses in animals (31, 85, 86), in order to account for the decreased overall synteny conservation among Brassicaceae. Furthermore, the syntenic alignments were screened for the presence of enabler substitutions, represented by a novel start codon, the removal of stop codons and/or frameshifts in the DNG coding region compared to the syntenic DNA of the outgroup species. Overall, we found 604 *A. thaliana* putative DNGs with apparent synteny conservation with at least one other Brassicaceae genome.

### Identification of *Brassica rapa* de novo genes

We retrieved all *Brassica rapa* genome assemblies and gene sets available as of October 2021 and screened each assembly for completeness using BUSCO v3.0 (87) (**S3 Table**). The *B. rapa* coding regions, protein and genome assembly fasta files were downloaded from: https://ngdc.cncb.ac.cn/search/?dbId=gwh&q=GWHAAES00000000.

*B. rapa* proteins containing stops (“Xs”) within their amino acid sequences were removed, leaving 45,912 proteins. We further screened the remaining *B. rapa* proteins to identify and remove sequences mostly formed by transposable elements (TEs), as they likely represent misannotated TEs. To this aim, we first downloaded the sequences of 39,197 TE families from 31 Brassicaceae reported in PlantRep (88). Proteins containing TE sequences were retrieved by performing a tBlastn search with the following modified parameters: -evalue 1e-10 - max_target_seqs 10 -max_hsps 5.

A total of 9,791 *B. rapa* proteins shared sequence similarity with TEs over at least 50% of their length and were removed from the dataset. Proteins with Blast matches uniquely with unknown repeats were not removed as those repeats might represent microsatellites, which could potentially form a portion of de novo gene coding regions.

To search for DNGs among the remaining 36,121 *B. rapa* proteins, we carried out a multipronged homology search strategy to identify proteins with homology in *non-Brassica rapa* genomes. First, the *B. rapa* proteins were searched against the plant NCBI refseq protein set obtained on August 31, 2021 throughout a Blastp run with the following modified parameters: - num_descriptions 5 -num_alignments 5 -evalue 0.001 -seg yes. A total of 18,413 proteins showed no sequence homology to NCBI refseq proteins with the exception of *Brassica* sequences, thus representing candidate *Brassica* orphan proteins. A further Blastp search was performed against the NCBI non-random, tsa_nr, refseq_rna and est databases of all Brassicaceae proteins with the following modified parameters: -max_target_seqs 50 -max_hsps 5. All hits containing premature stop codons were removed as they could represent expressed sequences of noncoding genes or truncated and thus non-functional proteins.

Similarly to the procedure applied to *A. thaliana* putative DNGs, we carried out Blast searches against increasingly broad taxonomic units containing *B. rapa*, starting from *Brassica* but excluding *B. rapa*, then other *Brassicaceae*, rosidae, angiosperms, green plants, excluding the previous taxon at each step, and using the following modified parameters: -max_target_seqs 50 - max_hsps 5-evalue 0.001 -seg yes. Fungal proteomes were also screened, whereas Archaea and Bacteria sequences were not included as our analyses in *A. thaliana* DNGs showed that prokaryotic databases contributed marginally to the detection of homologs. We found 2,089 *B. rapa* proteins sharing no sequenced homology with two or more non-*Brassica* proteins, thus representing *B. rapa* orphan genes. A cut-off of at least two non-*Brassica* proteins was implemented to take into account possible contamination from *A. thaliana* into other Brassicaceae genome datasets (89, 90). This number of orphan genes in *B. rapa* is similar to but higher than the 1,540 *B. rapa* orphan genes recently described (45), probably because of differences in the homology search criteria and in the gene annotation version. We further removed from the list of orphan genes 35 genes with annotated protein domains in eggNOG Mapper (91), as they likely represent fast evolving non-DNGs.

In order to identify enabler changes uniquely associated with de novo gene birth, we inspected the alignments from the Blast searches between *B. rapa* orphan genes and the genome of 45 Brassicaceae (**S2 Table**). After applying the same approach described to filter *A. thaliana* putative DNGs, we obtained 754 *B. rapa*-specific de novo genes.

### Examination of DNG coding and protein sequence features

#### DNA sequence features

Length of genes, predicted coding sequences (CDS) and (where available) UTRs were retrieved from gff files of the assemblies of each species. The GC-content of each coding region was calculated using an in-house perl script. Transposable element (TE) genome coordinates were obtained from the TAIR10 gff3 file in the *A. thaliana* TAIR repository and from the Brapa_genome_v3.0_TE.gff file downloaded from http://brassicadb.cn/#/Download/ for *B. rapa*.

*O. sativa* de novo gene IDs were retrieved from supplementary information in Zhang et al. (8). *O. sativa* release v3 coding region fasta sequences were downloaded from the Gramene repository (http://ftp.gramene.org/oge/release-3/fasta/oryza_sativa/dna/). Rice protein fasta sequences were obtained translating the CDS sequences using the ORF finder program in the SMS suite (92). The initial set of 50,556 genes was parsed to retain only the longest isoform of each locus and to remove genes with premature stop codons, leaving 38,748 genes.

Microsatellites were retrieved from the coding regions of each gene using Tandem Repeat Finder v4.09 (93) with default options. Overlap of TEs and microsatellites with the coding region and gene distance from TEs were obtained using bedtools (94). The coding potential was estimated using the Coding Potential Calculator (72). The codon adaptation index was calculated using the ‘cai’ tool in the EMBOSS suite (95).

Kozak scores were computed as the sum of the logs of nucleotide probabilities within a window of 12 bp around the ATG start codon (96), based on probabilities extracted from all genes in each species genome:

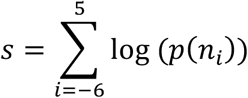

where *n_i_* is the observed nucleotide at position *i* relative to the start of the ATG. The scores are generally negative with a mean around −5.1, but the closer to zero, the more like the consensus sequence (AAAAAAATGGCG) they are. This is similar, but not identical, to the acAACAATGGC consensus sequence for terrestrial plants (97), reflecting biases that promote translation initiation by the ribosome. The nucleotide probability profiles surrounding start codons in *A. thaliana, B. rapa*, and *O. sativa*, as well as other diverse plant genomes (*Petunia inflata*: 36,489 genes; *Quercus robur*: 25,808 genes) are highly similar, although genes in *O. sativa* appear to have a relaxed constraint in the nucleotide following the ATG, whereas it is guanine over 50% of the time in the other two species (see **S1 Fig**), which could be related to fact that rice is a monocot and thus distantly related to the dicot family Brassicaceae. The preference for adenines upstream of the ATG in plants is much less pronounced than in nucleotide profiles of Kozak sequences in other eukaryotes (e.g. human, *Drosophila*) (see **S1 Fig**).

#### Protein sequence features

Protein domains were obtained from the NCBI Conserved Domain Database (98). Protein structural disorder was calculated using IUPred2 (99) after removing cysteines from the protein sequences in order to account for the possible presence of the disulfide bonds, which can strongly affect ISD estimates (100). Transmembrane helices were estimated using TMHMM Server v. 2.0 (101). The identified helices may represent either transmembrane structures or signal peptides, which tend to occur within the first 60 amino acid in the N-terminal of the protein. Proteins were conservatively assigned putative transmembrane helices if they contained one or more predicted helices and at least 18 amino acids in a helix past the first 60 amino acids. Signal peptides were predicted using TargetP2.0 (102). The isoelectric point of each protein was calculated using the Sequence Manipulation Suite Protein Isoelectric Point tool (92). All programs above were run using default settings.

*Functional genomic features in* A. thaliana. The *A. thaliana* TAIR10 gff3 file “TAIR10_GFF3_genes.gff” in the TAIR repository was used to obtain the length of “5’UTR length” and “3’UTR length” and the number of coding exons (“#Exons”). The proportion of the coding region overlapping with transposable elements (TEs; “TEs in CDS (bp)” and “%TEs in CDS”) and the gene distance to the nearest TE (“TEdist”) were calculated using bedtools (94) and the genome coordinate of TAIR10 coding exons and TEs. Genome coordinate of TEs were obtained from the gff3 file “TAIR10_GFF3_genes_transposons.gff” available in the TAIR repository. Possible regulatory motifs (“#Motifs promoter”) in the promoter regions of DNGs and AGs were identified using the MEME suite (103). The DNA sequences corresponding to the 300bp upstream of the transcription start site of each TAIR10 gene were retrieved using bedtools and screened using the MEME Streme tools (104).

Transcription factor binding site (TFBS) information was retrieved from the Plant *cis*-Map genome browser (http://ucsc.gao-lab.org/index.html). The Conserved TFBS dataset included binding sites deposited in the PlantRegMap database (105, 106). Conservation of TFBSs was assessed using multiple genome alignments across species from the Plant *cis*-Map genome browser conservation track. Binding sites with at least 50% of their sequence falling within conserved elements were considered conserved (“#Conserved TFBSs”). The pipeline to identify putative functional transcription factor binding sites (“#FUN TFBSs”) is described in Tian et al. (105). The AtRegNet confirmed TFBSs dataset (“#AtRegNet TFBSs”) was downloaded from the AGRIS database (107). The bedtools suite was used to extract the DNA sequences of the 200bp upstream of transcription start site of TAIR10 genes, corresponding to the putative promoter regions, and intersected with TFBS genome coordinates.

Transcription quantification features were obtained from a study of 18 natural *A. thaliana* accessions (108) study. The average (“RNAseq AVG”) and maximum (“RNAseq MAX”) expression across 48 samples, reported as log(rpkm) values, were calculated for each gene across 45 samples. Expression in only 1 out of 45 samples was also added (“RNAseq <2 samples”). Average (“RP AVG”) and maximum (“RP MAX”) ribosome profiling (RP) expression data, reported as log(rpkm) values, were also calculated for root tissues including control and deficient phosphorous nutrition conditions in a total of 25 samples (109). The maximum RP expression was calculated only for genes expressed in at least two samples.

The “Missense variant” feature represents the number of missense (nonsynonymous) substitutions divided by the length of the coding in each gene (“Missense variation”). Missense substitutions were obtained from the 1001 *A. thaliana* Genomes portal (https://1001genomes.org/index.html).

Protein domains and protein-protein interactions (PPIs) were obtained from the files TAIR10_all.domains (“#HMMPfam Domain”) and TairProteinInteraction 20090527.txt (“#PPIs”, “#PPIs (w/predicted)”), respectively, from the TAIR repository. The PPI data contain interaction annotations extracted from the literature by TAIR and BIOGRID (110). The frequency of three different categories of amino acids, “Tiny”, “Aromatic” and “Acidic”, were obtained using the pepstats program in the EMBOSS suite (95).

We gathered phenotypic information using data deposited in the TAIR repository containing phenotypic data extracted from the literature by TAIR. Data files names: TAIR_Phenotypes_9- 2019.txt (“TAIR_Phenotypes_9-2019.txt”), Locus_Germplasm_Phenotype_20190630.txt (“LGP_20190630.txt”), Locus_Germplasm_Phenotype_20130122 (“LGP_20130122”). Additionally, data from manually curated meta-analysis of loss-of-function phenotypes (111) (“Lloyd and Meinke 2012”) and from a high-throughput phenotype screening of annotated genes with unknown function (112) (“Luhua, et al. 2013”) were included.

### Machine learning training and testing

Balanced sets of DNGs and AGs were selected for ML training and testing within each species. For instance, in *A. thaliana* 331 AGs were randomly chosen among the 26,423 available AGs. The Decision Tree (DT) classifier was trained using the *scikit-learn* package in Python (113). The feed-forward fully-connected neural network (NN) with a single hidden layer with 20 hidden nodes was also trained with the MLPClassifier implementation in *scikit-learn*, using *tanh* activation functions and the ‘adam’ solver. Log transformations were applied to length-based features (Gene length, CDS length, Distance from TEs) and to functional genomic features with RPKM in *A. thaliana*.

Feature importance (normalized decrease in Gini impurity index at each node where a feature is used, weighted by the fraction of training examples represented at those nodes) was calculated using the ‘feature_importance’ attribute of the DecisionTreeClassifier generated by *scikit-learn*. In some runs, “At-least-one-domain” was the feature at the root of the tree, and in other runs, “Coding potential (cpc2)” represented the splitting feature at the root. Thus, we estimated the importance of features by averaging them over multiple runs.

Scripts and data from this study are available at https://github.com/ioerger2/DNG. This repository contains instructions on how to run Decision Tree and Neural Network training and testing. A python script generates accuracies, confusion matrices, decision trees and feature importances for each dataset. The *A. thaliana* features are available in the repository.

## Statistical analyses

All statistical analyses were performed in R (cit). In MLA testing, accuracy corresponds to the sum of true positives (TPs) and true negatives (TNs), divided by all genes: (TP+TN)/(TP+FP+TN+FN). Sensitivity and specificity are represented by true positive rate (TPR=TP/(TP+FN)) and true negative rate (TNR=TN/(TN+FP)). Test sets and training sets are always kept disjoint (with no overlap of genes) in all runs.

## Acknowledgements

The authors would like to thank Michael Dickens and the Texas A&M University High Performance Research Computing facility for assistance with performing homology search analyses on HPC clusters.

## Author Contributions

CC, AEP and TRI conceived and designed the study. CC, AO and TRI analyzed the data. TRI.

CC, AEP and TRI wrote the paper.

