## Supplementary material for "Accurate identification of de novo genes in plant genomes using machine learning algorithms": S3 Fig

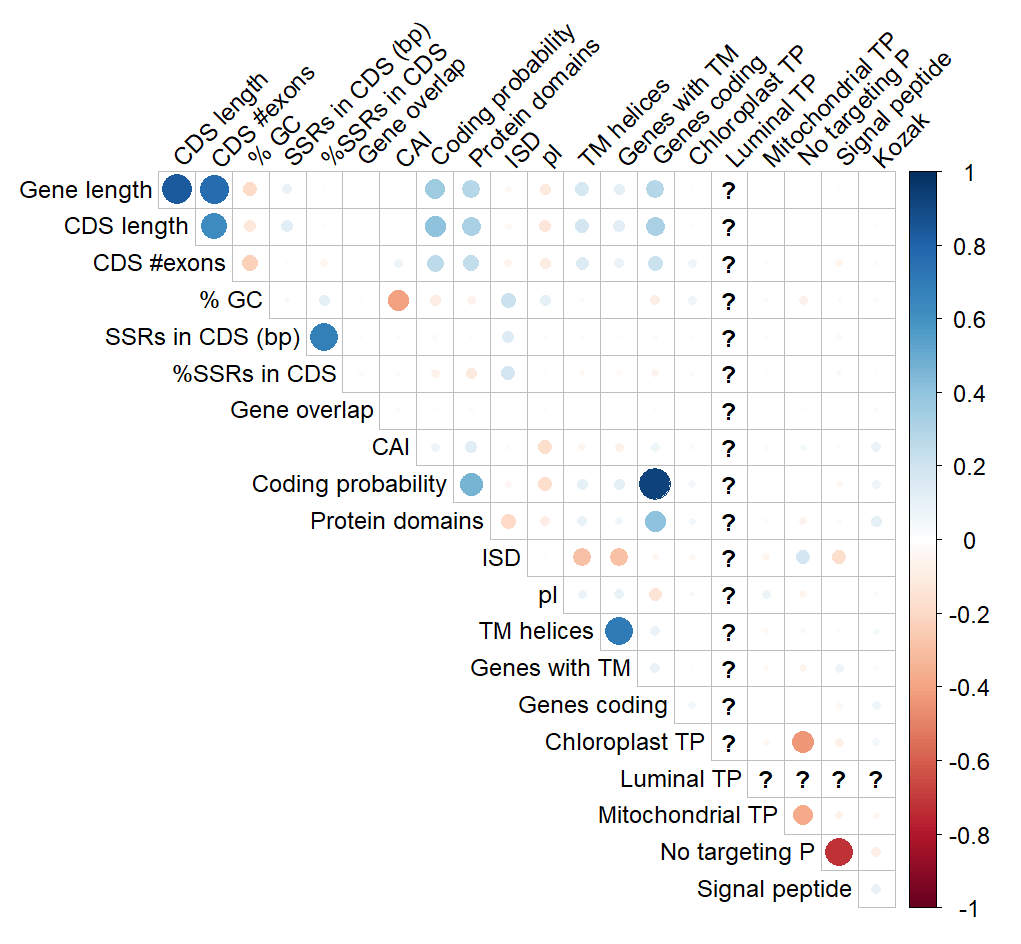

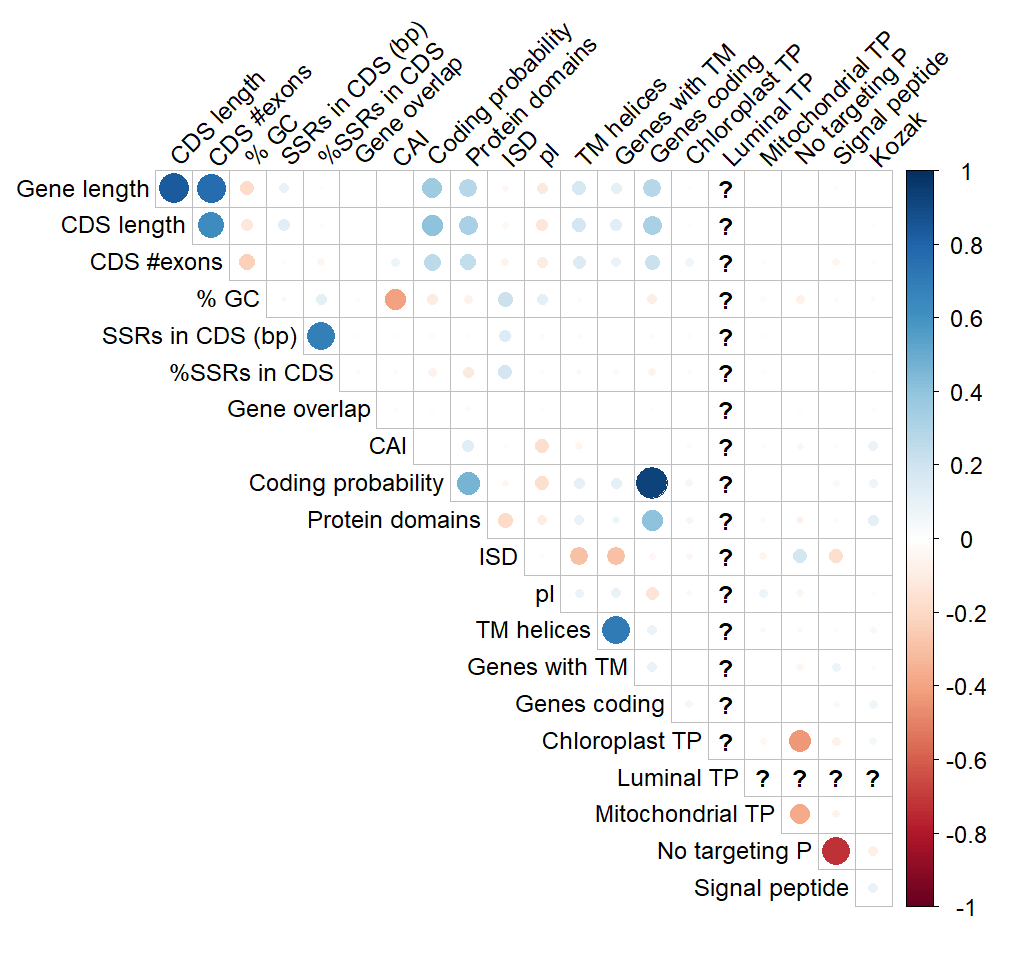

**S3 Fig**. **Feature correlations for *B. rapa* (left) and *O. sativa* (right).** Note: the ?’s in the row and column for ‘Luminal TP’ is because this feature was 0 for all genes in *B. rapa* *and O. sativa*, hence covariances could not be calculated.
