## Supplementary material for "Accurate identification of de novo genes in plant genomes using machine learning algorithms": S2 Fig

A
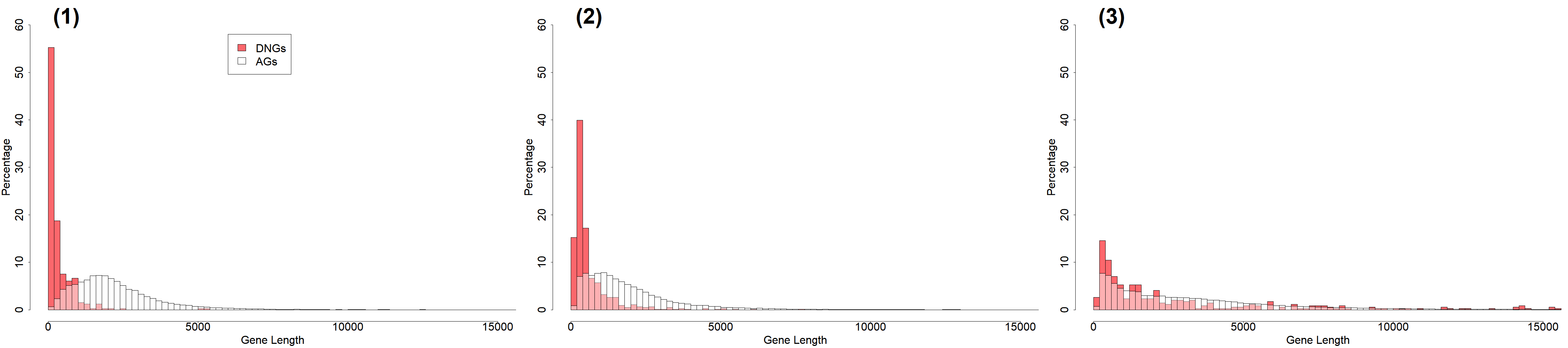

B
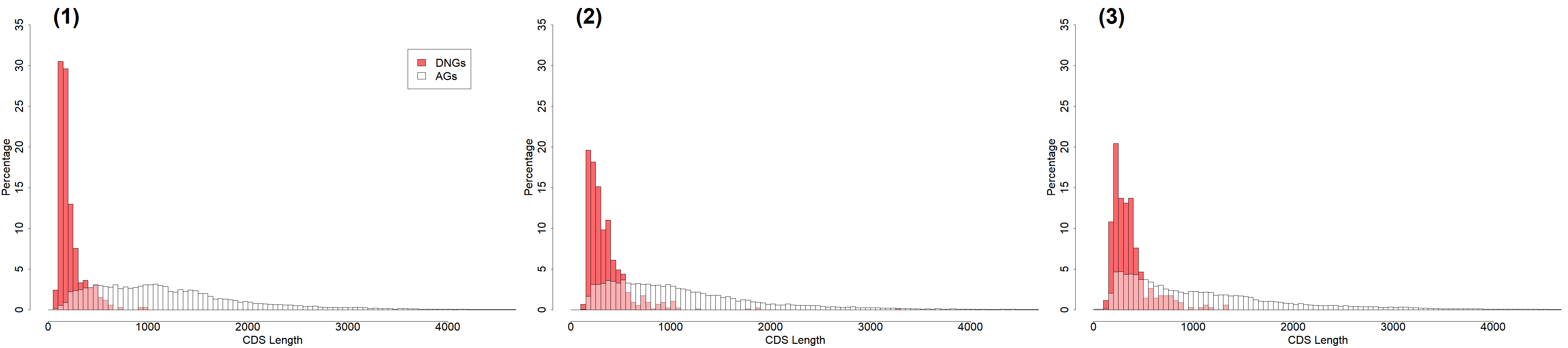

C
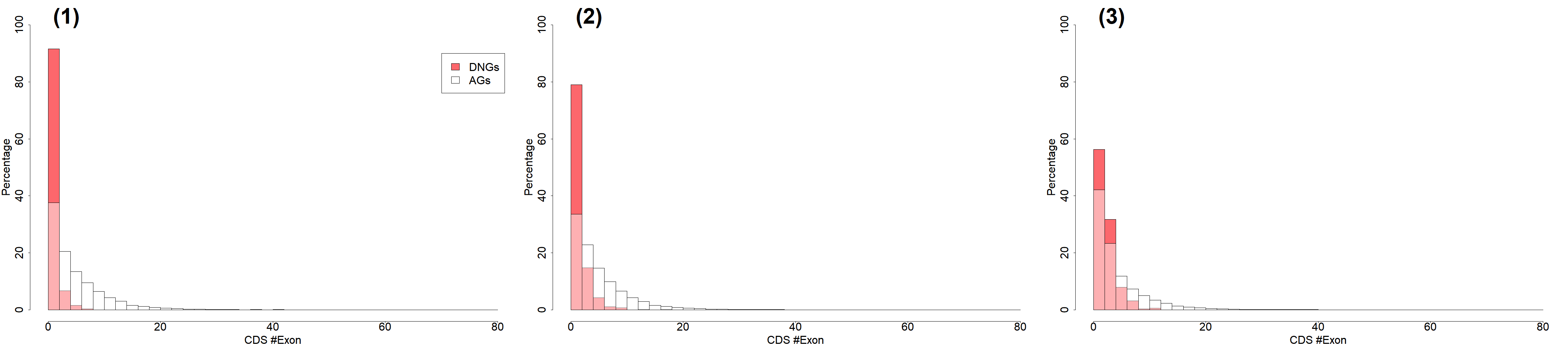

D

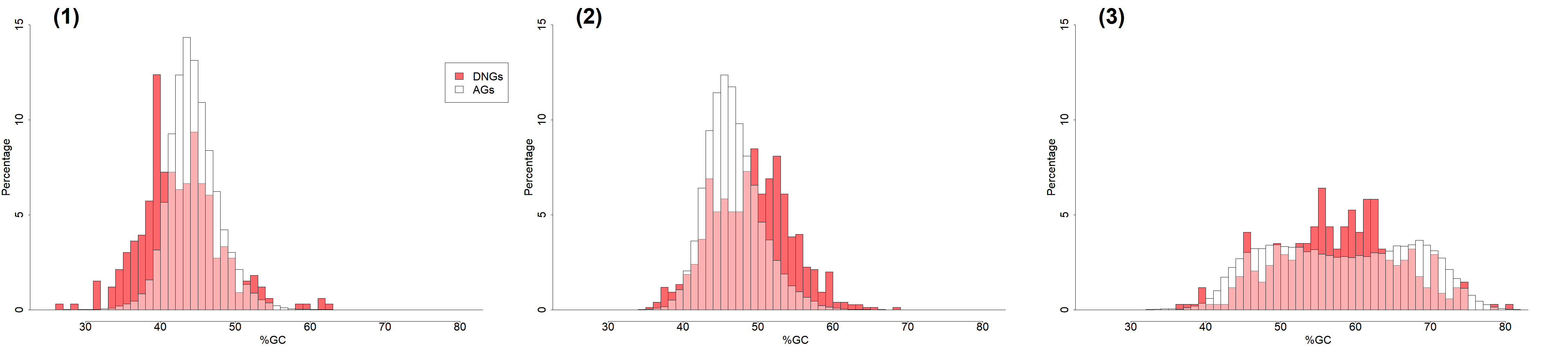

E
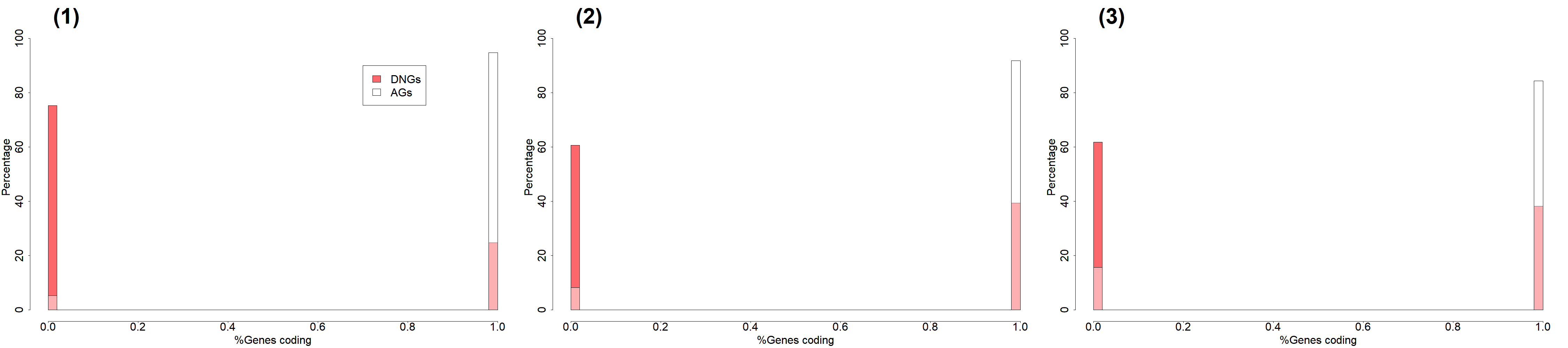

F
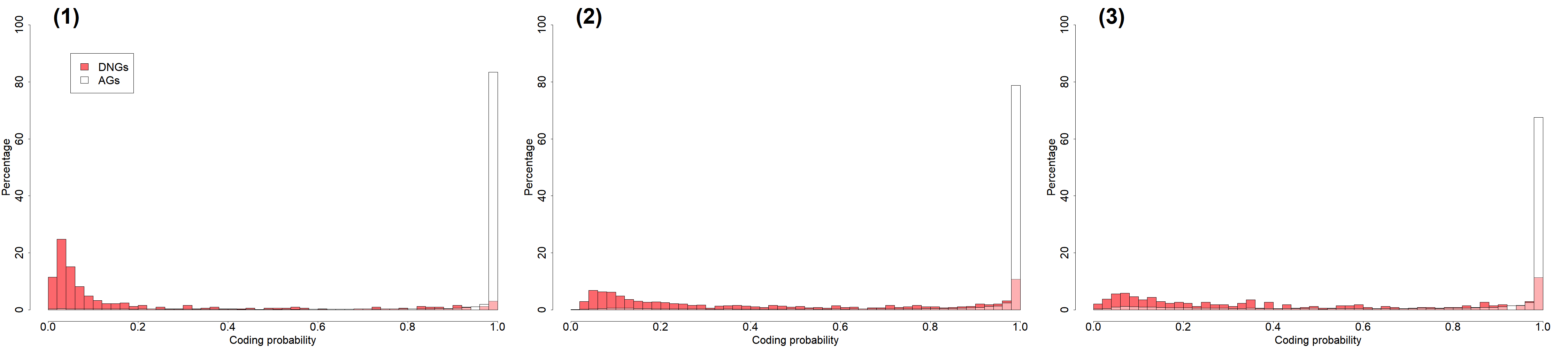

G
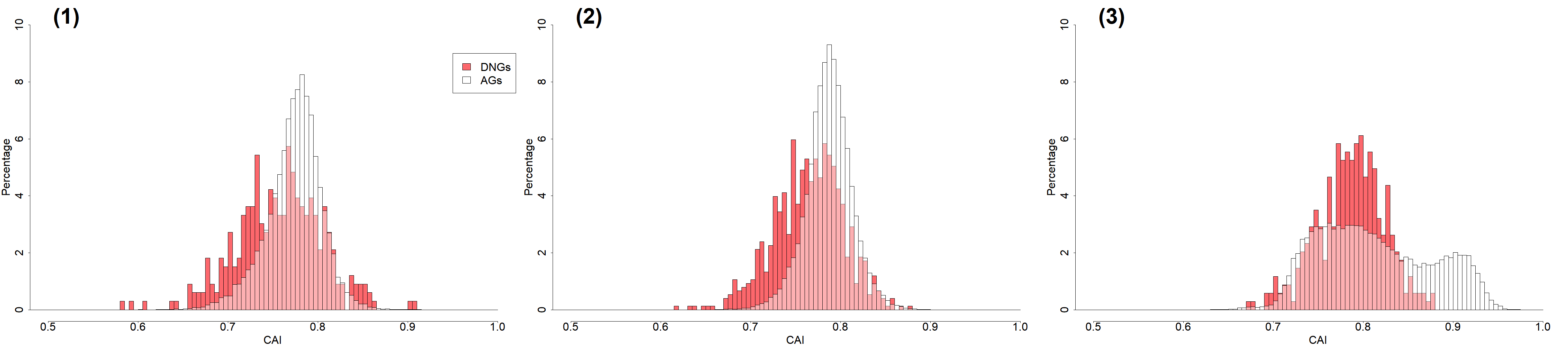

H
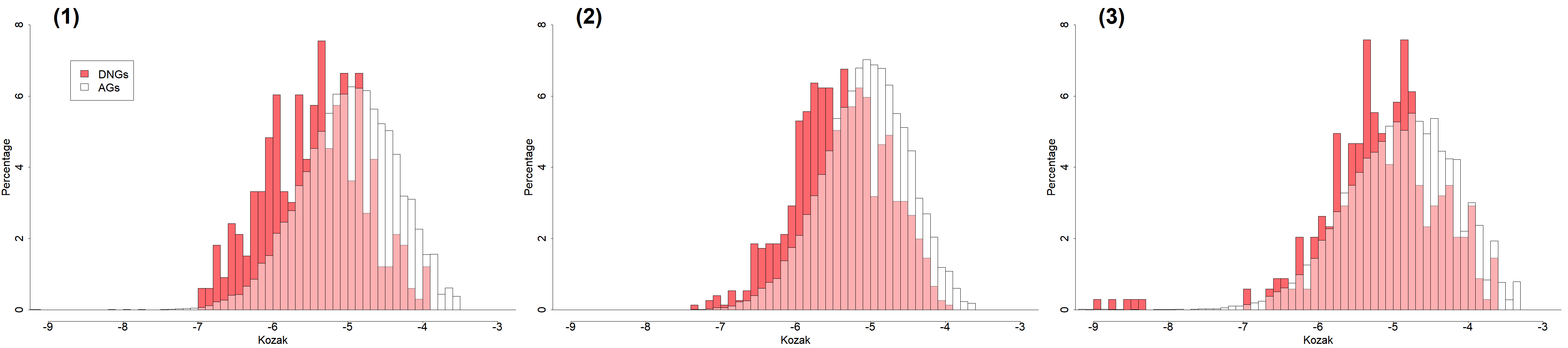

I
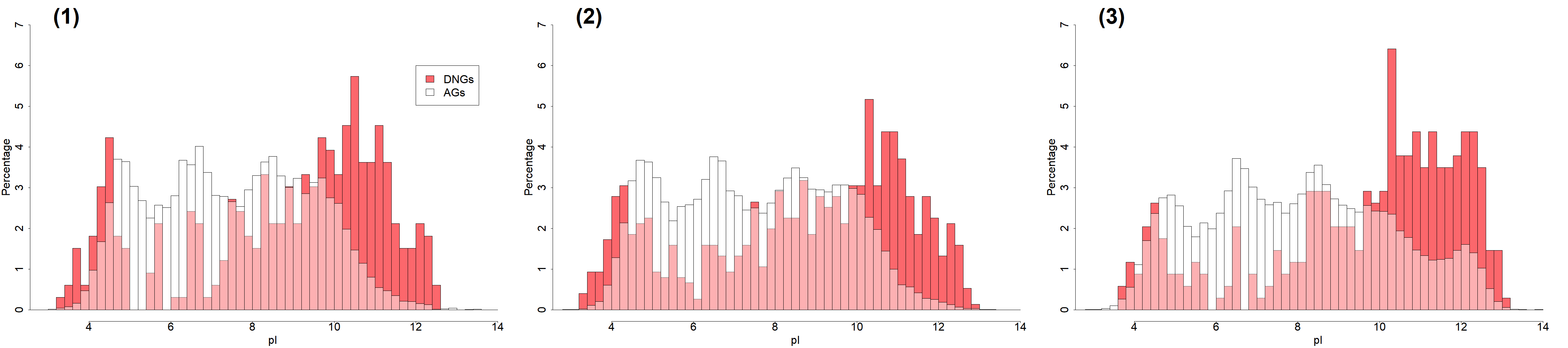

K
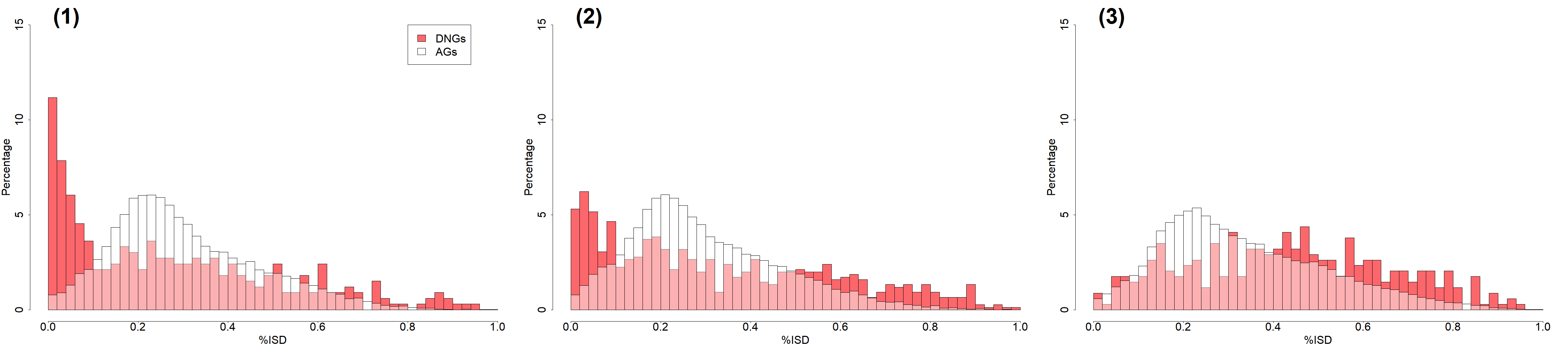

**S2 Fig**. Distribution of values for major sequence features in DNGs and AGs. Left panels (1): *A. thaliana*. Middle panels (2): *B. rapa*. Right panels (3): *O. sativa*.
