## Supplementary material for "Accurate identification of de novo genes in plant genomes using machine learning algorithms": S1 Fig

| *Arabidopsis thaliana*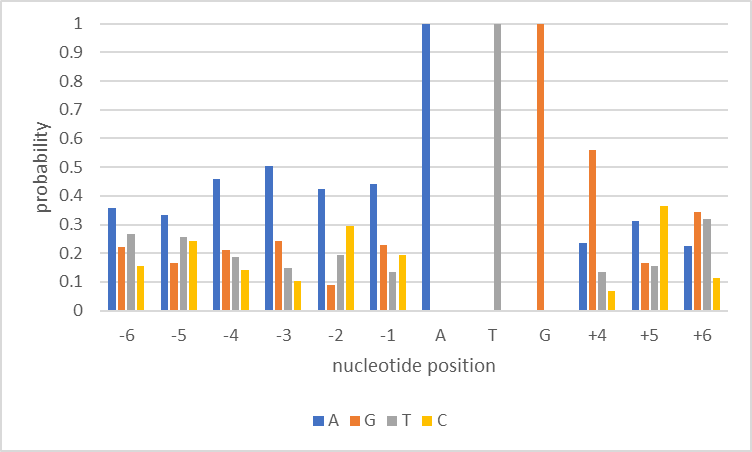 | *Brassica rapa*  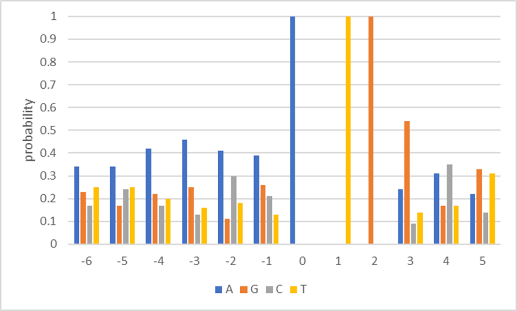 | *Oryza sativa japonica*  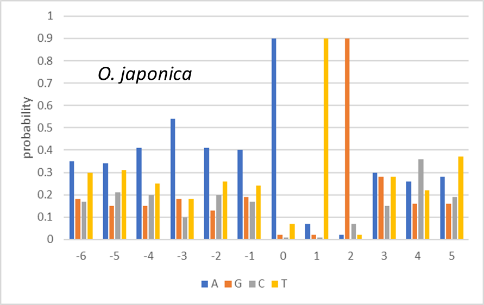 |
| --- | --- | --- |
| 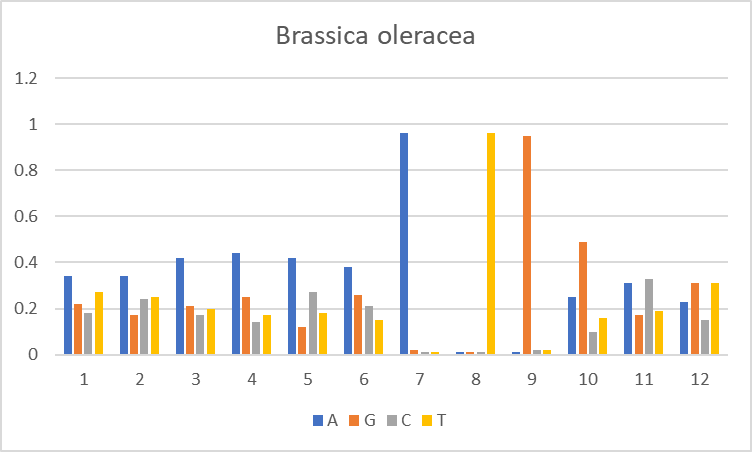 | 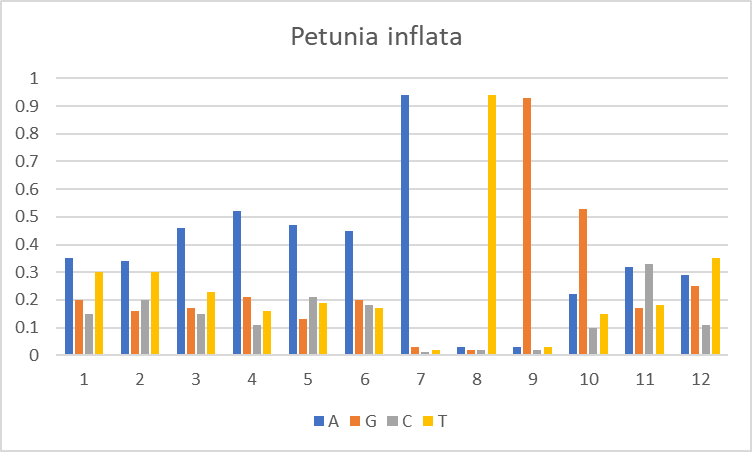 | 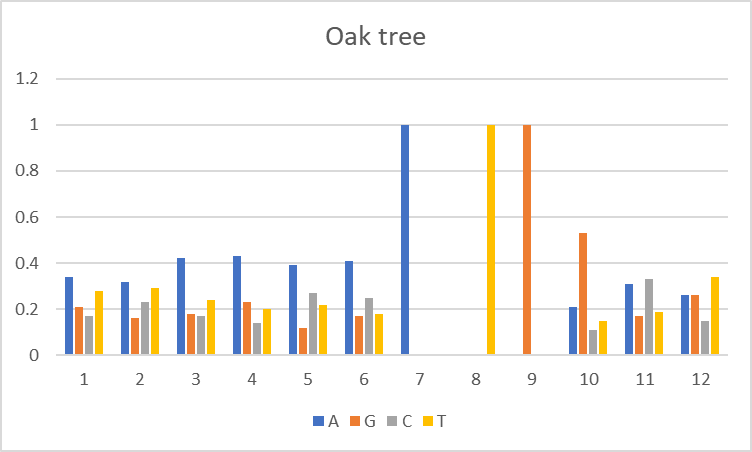 |
| *Saccharomyces cerevisiae*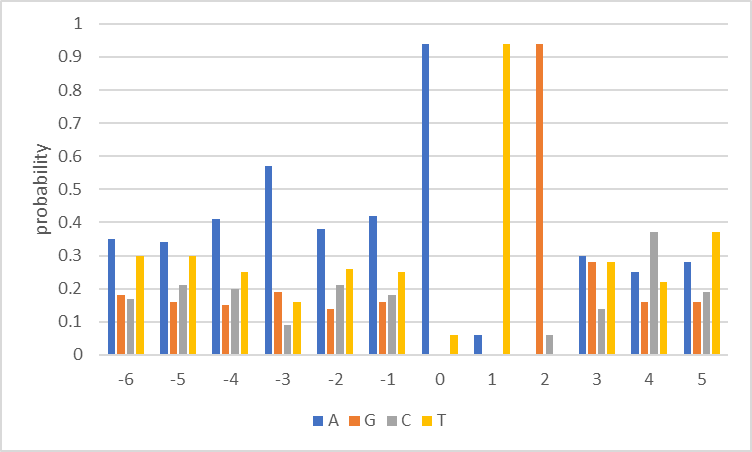 | *Drosophila melanogaster*  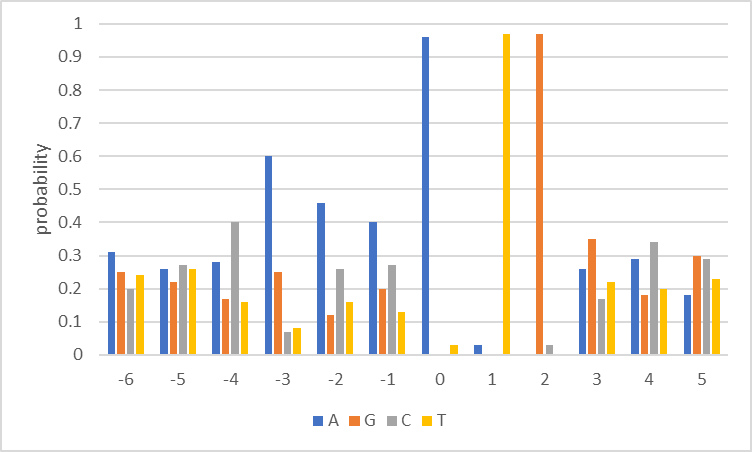 | Human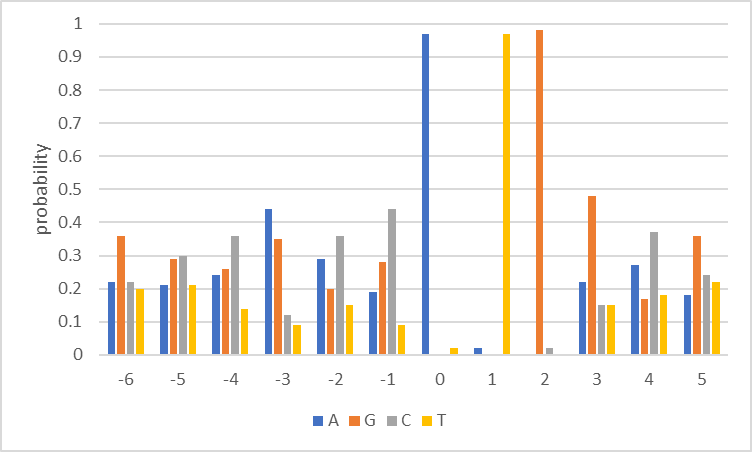 |

**S1 Fig**. Kozak Sequence Profiles. Nucleotide probability profiles in a 12 bp window around the start codon for all CDS annotated in a genome. Note that the “ATG” is apparent at positions 0-2. Notice the preference for ‘A’ at all 6 positions upstream of the ATG in plant genomes. Furthermore, ‘G’ is preferred the nucleotide immediately after the ATG (except for in the rice genome, *O. sativa*). These features are not as apparent in other eukaryotes.
